# Wss1 promotes replication stress tolerance by degrading histones

**DOI:** 10.1101/2019.12.13.876110

**Authors:** Karthik Maddi, Daniel Kwesi Sam, Florian Bonn, Stefan Prgomet, Eric Tulowetzke, Masato Akutsu, Jaime Lopez-Mosqueda, Ivan Dikic

**Affiliations:** Institute of Biochemistry II, Goethe University School of Medicine, Theodor-Stern-Kai 7, 60590 Frankfurt am Main, Germany; Buchmann Institute for Molecular Life Sciences, Goethe University, Max-von-Laue-Str. 15, 60438 Frankfurt am Main, Germany; South Dakota State University, Department of Biology and Microbiology, Brookings South Dakota, USA

**Keywords:** Wss1, metalloprotease, Hydroxyurea (HU), histone H3, H2A and H4

## Abstract

Timely completion of DNA replication is central to accurate cell division and to the maintenance of genomic stability. However, certain DNA-protein interactions can physically impede DNA replication fork progression. Cells remove or bypass these physical impediments by different mechanisms to preserve DNA macromolecule integrity and genome stability. In *Saccharomyces cerevisiae*, Wss1, the DNA-protein crosslink repair protease, allows cells to tolerate hydroxyurea-induced replication stress but the underlying mechanism by which Wss1 promotes this function has remained unknown. Here we report that Wss1 provides cells tolerance to replication stress by directly degrading core histone subunits that non-specifically and non-covalently bind to single-stranded DNA. Unlike Wss1-dependent proteolysis of covalent DNA-protein crosslinks, proteolysis of histones does not require Cdc48 nor SUMO-binding activities. Wss1 thus acts as a multi-functional protease capable of targeting a broad range of covalent and non-covalent DNA-binding proteins to preserve genome stability during adverse conditions.

## Introduction

DNA-bound proteins can be physical impediments to replication forks and associated protein complexes. DNA-protein crosslinks (DPCs) in particular, constitute physical obstacles to DNA and RNA polymerases (Edenberg et al., 2014) and are genotoxic to cells because direct collisions between replication forks and DPCs covert DPCs into DNA double strand breaks. In budding yeast, Wss1 was identified as the first DNA-dependent protease important for DPC repair. Wss1-dependent DPC repair requires physical interactions with the ATPase Cdc48 (p97/VCP in mammals), and binding to the small ubiquitin-like modifier, Smt3 (SUMO in higher eukaryotes). Wss1 mutants deficient in binding to either Cdc48 or Smt3 fail to provide tolerance to DPC-inducing agents. In higher eukaryotes, SPRTN is the DNA-dependent protease, which removes proteins that become covalently attached to DNA (Lopez-Mosqueda et al., 2016; Stingele et al., 2016; Vaz et al., 2016). However, unlike Wss1, SPRTN can repair DPCs independently of its ability to bind to p97 (Cdc48) or Ubiquitin (Lopez-Mosqueda et al., 2016).

Replication fork barriers, such as DPCs, can be bypassed by several distinct mechanisms (Duxin et al., 2014; Sparks et al., 2019). Wss1, however, can directly and irreversibly remove the DPC protein component by proteolysis (Balakirev et al., 2015; Stingele et al., 2014). Wss1 is not essential for viability in budding yeast cells nor it is required for surviving chronic exposure to DPC-inducing agents including Camptothecin.

Mass-spectrometry based analysis of isolated DPCs identified DNA-binding proteins including replication proteins, transcription factors, and histones-histones constituting the most abundant protein components of DPCs (Vaz et al., 2016). Histones are abundant proteins that have a high affinity for DNA and their levels are carefully regulated at the transcriptional, translational and post-translational level. Owing to their positive charge, histones can form protein-DNA aggregates (Shintomi et al., 2005) that have been previously reported to impede gene transcription (Karsten, 2008). Proteins do not necessarily need to be covalently attached to DNA to constitute a replication fork barrier. To exemplify, Fob1 in yeast (Kobayashi and Horiuchi, 1996) and the viral protein Epstein-Barr nuclear antigen 1 (EBNA-1) in mammalian cells (Dhar and Schildkraut, 1991) bind to DNA with high affinity and their binding effectively block replication fork progression. Here we show that Wss1 is important for replication stress tolerance. Wss1 directly elicits proteolytic degradation of histones that are non-specifically bound to single-stranded DNA. Wss1-dependent replication stress tolerance, as well as histone degradation, do not require Cdc48 or Smt3 binding activities. Our results suggest that Wss1 is a multi-functional protease that irreversibly removes covalently and non-covalently bound proteins from DNA to provide replication stress tolerance.

## Results

### Wss1 proteolytic activity is necessary for replication stress tolerance

Wss1 is dispensable for yeast viability due, in part, to the redundant pathways that mediate DPC repair (Stingele et al., 2017; Vaz et al., 2017). However, cells deficient in Wss1 function display a strong negative genetic interaction with Tdp1, a tyrosyl-DNA phosphodiesterase that assists in the removal of DNA-protein crosslinks, such that cells lacking both, Wss1 and Tdp1, are synthetic lethal (Lopez-Mosqueda et al., 2016; Sharma et al., 2017; Stingele et al., 2014). Wss1 becomes essential for viability when DNA repair by homologous recombination is compromised, underscoring the notion that multiple pathways coalesce to repair DPCs (Stingele et al., 2014). To identify conditions that require Wss1 function, we screened various genotoxic agents in Wss1-deficient yeast cells (*wss1*Δ) and searched for a slow-growing phenotype (**Figure1A and Supplemental Figure 1A**). We observed that *wss1*Δ cells are sensitive to chronic exposure to the DNA replication inhibitor hydroxyurea (**Figure 1A**), which depletes deoxyribonucleotide pools by inhibiting ribonucleotide reductase (O’Neill et al., 2004) and are sensitive to hydrogen peroxide which produces reactive oxygen species (**Supplemental Figure Figure S1A**). Similarly, when *wss1*Δ cells are only transiently exposed to hydroxyurea, we observed a modest but reproducible viability loss compared to wildtype cells exposed to the same treatment (**Figure 1B**). While, it remains unknown whether hydroxyurea directly cause DNA-protein crosslinks, it is known that hydroxyurea is metabolized to form the free radical nitroxide(King, 2003; Yarbro, 1992). Free radicals are known DNA-protein crosslink inducers (Gajewski and Dizdaroglu, 1990; Nakano et al., 2003; Wickramaratne et al., 2016) raising the possibility that, at high concentrations, hydroxyurea may lead to DNA-protein crosslinks. Nevertheless, Wss1 plays an important role in providing cells tolerance to hydroxyurea-induced replication stress.

**Figure 1:**
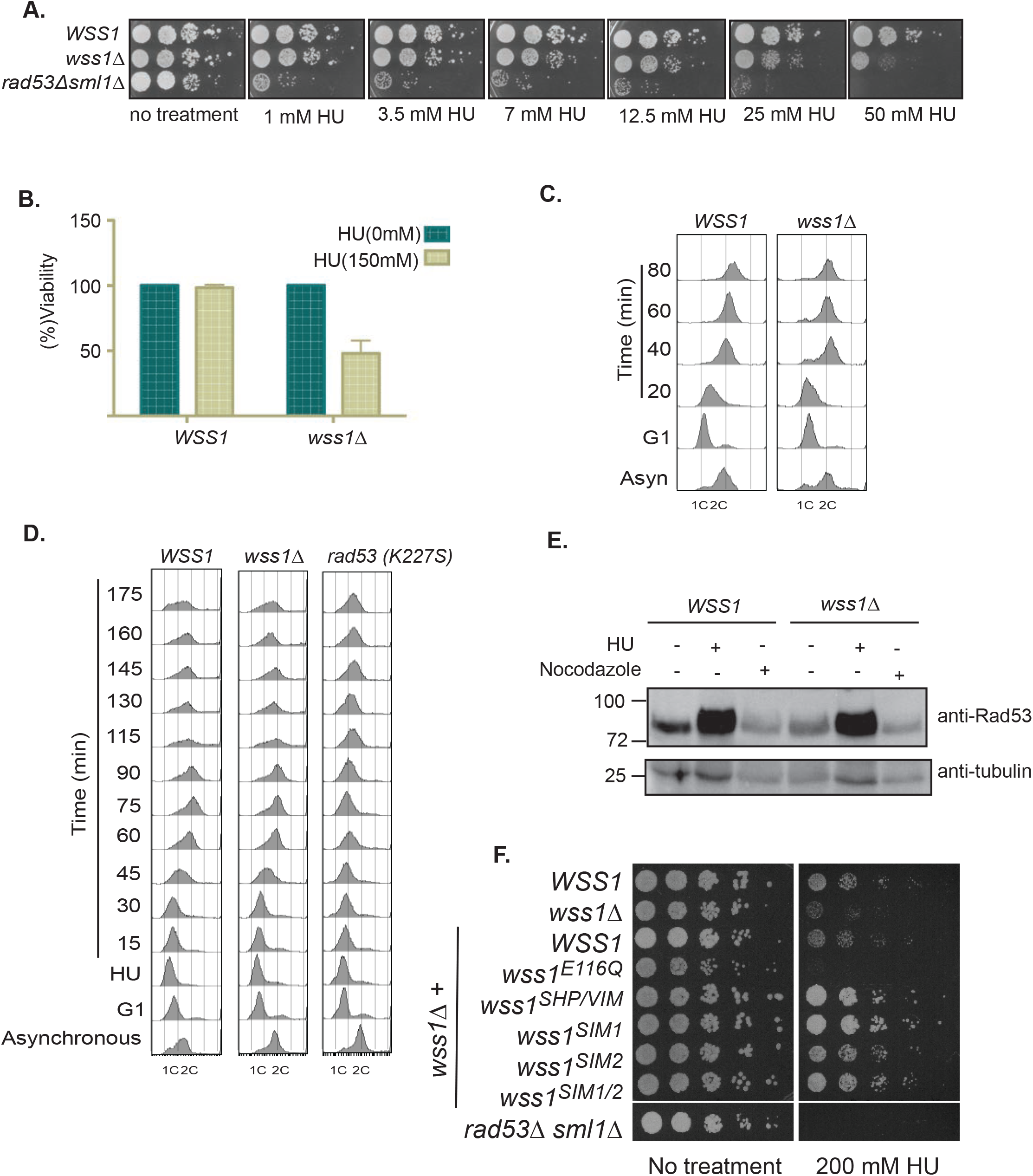
Wss1 proteolytic activity is necessary for replication stress tolerance. **A**) *wss1*Δ cells are sensitive to chronic exposure to hydroxyurea. Ten-fold dilutions of wild type, *wss1*Δ or *rad53*Δ cells were spotted on YPD plates in the absence or presence of hydroxyurea (1, 3.5, 7, 12.5, 25 and 50 mM) and incubated at 30°C. Plates were imaged after 3 days. **B**) *wss1*Δ cells are sensitive to transient exposure to hydroxyurea. Exponentially growing wildtype or *wss1*Δ cells were either untreated or treated with hydroxyurea (150 mM) for 1h and washed with YPD media to remove residual hydroxyurea. Approximately 2×10^3^ cells were plated on YPD plates and incubated at 30°C for 2 days. Colonies were counted using ImageJ and bars in the histogram represent percentage of viable cells after hydroxyurea treatment. **C**) Wss1 is not required for DNA replication fork progression. Flow cytometric (FACS) analysis of wildtype or *wss1*Δ cells synchronized in G1 with alpha-factor and released from the G1 arrest in the absence of hydroxyurea. **D**) Wss1 is not required for recovery after hydroxyurea. FACS analysis of wild type, *wss1Δ or rad53*Δ cells synchronously released from a G1 arrest into hydroxyurea (200 mM for 1 hr). Cells were washed and released in to YPD media without hydroxyurea and samples were collected at indicated time points. **E**) Wss1 is not required for Rad53 activation. Exponentially growing wildtype or *wss1*Δ cells were treated with hydroxyurea (200 mM) for 1 hour and whole cell extracts were prepared by TCA precipitation method and immuno-blotted using a Rad53 specific antibody or alpha-Tubulin for loading control. **F**) Wss1 domain requirements for hydroxyurea tolerance. Five-fold dilutions of *wss1*Δ cells reconstituted with wildtype or wss1 mutants and cells were spotted on plates with or without hydroxyurea (50, 100 and 200 mM) and incubated at 30°C. Plates with 50 and 100 mM hydroxyurea were imaged after 2 days whereas plates with 200mM hydroxyurea were imaged after 4 days.

To account for the hydroxyurea sensitivity observed in *wss1*Δ cells, we first asked whether *wss1*Δ cells have defects traversing S-phase. Wildtype (*WSS1*) and *wss1*Δ cells were arrested in the G1 phase of the cell cycle with alpha-factor and released from the G1 block into media without alpha-factor to allow cells to enter synchronously into the S-phase. We monitored DNA replication dynamics using flow cytometry. Under these conditions, we did not observe an appreciable defect in bulk DNA replication in *wss1*Δ cells as *wss1*Δ cells entered and progressed through S-phase with similar kinetics as wildtype cells (**Figure 1C**). We next asked if Wss1 is important for recovery from replication fork stalling. To achieve replication fork stalling, we first blocked cells in G1, then released cells into media containing hydroxyurea and held them for 1 hour. Finally, cells were released from the hydroxyurea treatment and we monitored replication dynamics using flow cytometry as before. The DNA damage checkpoint kinase, Rad53, is critical for maintaining replication fork stability in hydroxyurea (Desany et al., 1998). As such, *rad53*Δ cells are hypersensitive to hydroxyurea and do not recover from a hydroxyurea treatment. In contrast to *rad53*Δ cells, *wss1*Δ cells recovered from hydroxyurea similarly to wildtype cells (**Figure 1D**). We tested whether Wss1 is important for Rad53 activation, as failure to activate Rad53 would result in hydroxyurea sensitivity. We assayed for Rad53 auto-phosphorylation as a measure of Rad53 activation in wildtype and *wss1*Δ cells and did not observe a marked defect in Rad53 activation in *wss1*Δ cells exposed to hydroxyurea (**Figure 1E**). Taken together, the hydroxyurea sensitivity observed in *wss1*Δ cells is not due to major defects in completing DNA replication or a general failure to stably maintain replication forks.

To better understand the hydroxyurea induced toxicity observed in *wss1*Δ cells, we next investigated which Wss1 activity is needed to mediate hydroxyurea tolerance. Wss1 is endowed with the ability to physically interact with Cdc48 and Smt3 owing to its SHP/VIM and tandem SIM motifs, respectively. Wss1 is also able to degrade Top1 covalent complexes via its protease domain (SprT) (Balakirev et al., 2015; Stingele et al., 2014). We reconstituted *wss1*Δ cells with either wildtype Wss1 or mutant alleles deficient in protease activity (E116Q), Cdc48 binding (SHP & VIM) or Smt3 binding (SIM1, SIM2) and confirmed expression of wildtype and mutant Wss1 alleles by Western blot analysis (**Supplemental Figure 1C**). As expected, reconstitution of *wss1*Δ cells with wildtype Wss1 restored hydroxyurea resistance, whereas reconstitution with the catalytically inactive protease mutant did not (**Figure 1F**). Interestingly, reconstitution of *wss1*Δ cells with SHP/VIM and SIM mutant alleles also restores hydroxyurea resistance, suggesting that Cdc48 and Smt3 binding to Wss1 are dispensable to mediate hydroxyurea resistance (**Figure 1F**). Over-expressing *wss1-E116Q* mutant confers a lethal phenotype in hydroxyurea treated cells compared to cells where Wss1 is absent. This observation suggests that protein complexes that include Wss1 are accumulating in the presence of hydroxyurea. Taken together, Wss1 can target proteins for degradation to mediate hydroxyurea tolerance and this activity is independent of Cdc48 and Smt3 binding.

### Proteolytic targets of Wss1 upon hydroxyurea stress

Having established that the proteolytic activity of Wss1 is essential to overcome the hydroxyurea stress, we next sought to identify Wss1 target proteins, whose degradation would be important in mediating tolerance to hydroxyurea. We reasoned that putative Wss1 substrates would be more abundant in *wss1*Δ cells than in wildtype cells exposed to hydroxyurea. To quantitatively determine differences in protein abundance wildtype and *wss1*Δ were labeled in light and heavy Lysine, respectively using stable isotope labeling by amino acids in cell culture (SILAC). Whole cell extracts from SILAC-labeled wildtype and *wss1*Δ cells, under basal conditions and after hydroxyurea treatment, were generated from three biological replicates (**Figure 2A**) and processed for mass spectrometry. We identified 3,793 unique proteins representing ~ 84% of the expressed yeast proteome in the untreated and hydroxyurea-treated cells. Of the 3,793 proteins identified, 1,077 were up-regulated specifically in *wss1*Δ cells treated with hydroxyurea compared to 317 up-regulated proteins in wildtype cells treated with hydroxyurea (**Figure 2B**), which is consistent with our hypothesis that proteins would accumulate in *wss1*Δ cells. Using GO enrichment analysis, we observed an enrichment of proteins involved in regulation of DNA and RNA metabolism as up-regulated proteins in hydroxyurea-treated *wss1*Δ cells (**Figure 2C**). Our attention was next focused on three candidates - Rad51, Ddi1 and histone H3 - that were more abundant in hydroxyurea-treated *wss1*Δ cells (**Figure 2D and Supplemental Figure 2A**). In order to validate mass spectrometry-based results, Rad51 and Ddi1 were tagged at their carboxyl-terminal ends with three tandem copies of the FLAG epitope in wildtype and *wss1*Δ cells. Rad51 is a single-stranded DNA binding protein that is important for initiating recombination, while Ddi1 is an aspartic protease that functions as a shuttle factor for the proteasome. Rad51 accumulation on single-stranded DNA is toxic to cells and its accumulation is counteracted by Srs2 and BLM helicases. Yeast cells lacking Wss1 have previously been reported to have a hyper-recombination phenotype (Munoz-Galvan et al., 2013; Stingele et al., 2016). Western blot analysis of Rad51 protein levels in wildtype and *wss1*Δ cells reveals that Rad51 is more abundant in *wss1*Δ cells (**Figure 3A**). We reasoned that if Rad51 were to accumulate in *wss1*Δ cells during a hydroxyurea treatment and this Rad51 accumulation is the underlying reason for the hydroxyurea sensitivity, then the genetic ablation of Rad51 should suppress the hydroxyurea sensitivity observed in *wss1*Δ cells. To directly test this hypothesis, we generated *rad51*Δ and *wss1*Δ *rad51*Δ cells and found that *wss1*Δ *rad51*Δ cells were more sensitive to hydroxyurea than single mutants alones (**Figure 3B**). Similarly, Ddi1 was found to be upregulated in cells devoid of Wss1, as well as in cells treated with hydroxyurea (**Supplemental Figure 2A** **and** **2B**). We generated *ddi1*Δ and *wss1*Δ *ddi1*Δ cells and tested their abilities to tolerate hydroxyurea. *Ddi1*Δ cells alone were more sensitive to hydroxyurea than *wss1*Δ cells and a synergistic effect is observed in *wss1*Δ *ddi1*Δ cells (**Supplemental Figure 2C**).

**Figure 2:**
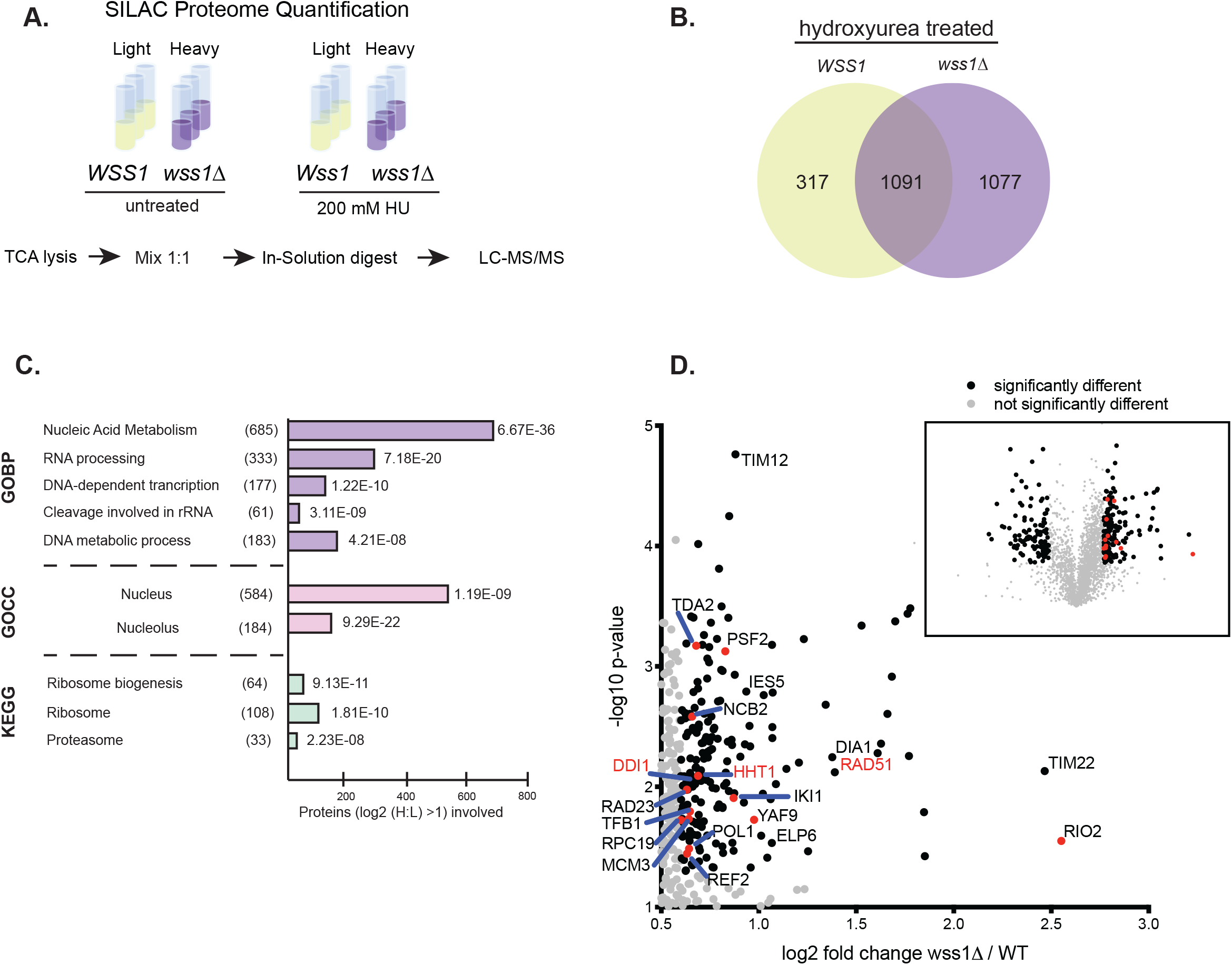
Excess histones are toxic to Wss1 mutant cells. **A**) Mass spectrometry-based proteome quantification. Experimental scheme used for quantifying protein abundance in wildtype or *wss1*Δ cells treated with or without hydroxyurea (200 mM). Cells were SILAC labelled with light or heavy Lysine. TCA extracts were generated after hydroxyurea treatment and mixed 1:1 prior to in-solution digest and subsequent analysis with LC-MS/MS. **B**) Venn diagram regulated protein identified in wildtype or *wss1*Δ cells treated with hydroxyurea. **C**) GO enrichment analysis. Proteins found to be statistically significantly enriched in *wss1*Δ cells to predict the molecular function of Wss1 in the presence of hydroxyurea. **D**) Volcano plots representing SILAC based quantification of peptides from Wild type and *wss1*Δ cells treated with hydroxyurea. Inset represents the results of three independent mass spectrometry based quantification of proteome changes in wildtype (left) and *wss1*Δ cells (right) treated with hydroxyurea. Gray circles represent proteins that were not statistically enriched. Black circles represent proteins that were significantly enriched *t-test p<0.5*. Protein labeled in red were confirmed by western blot analysis.

**Figure 3:**
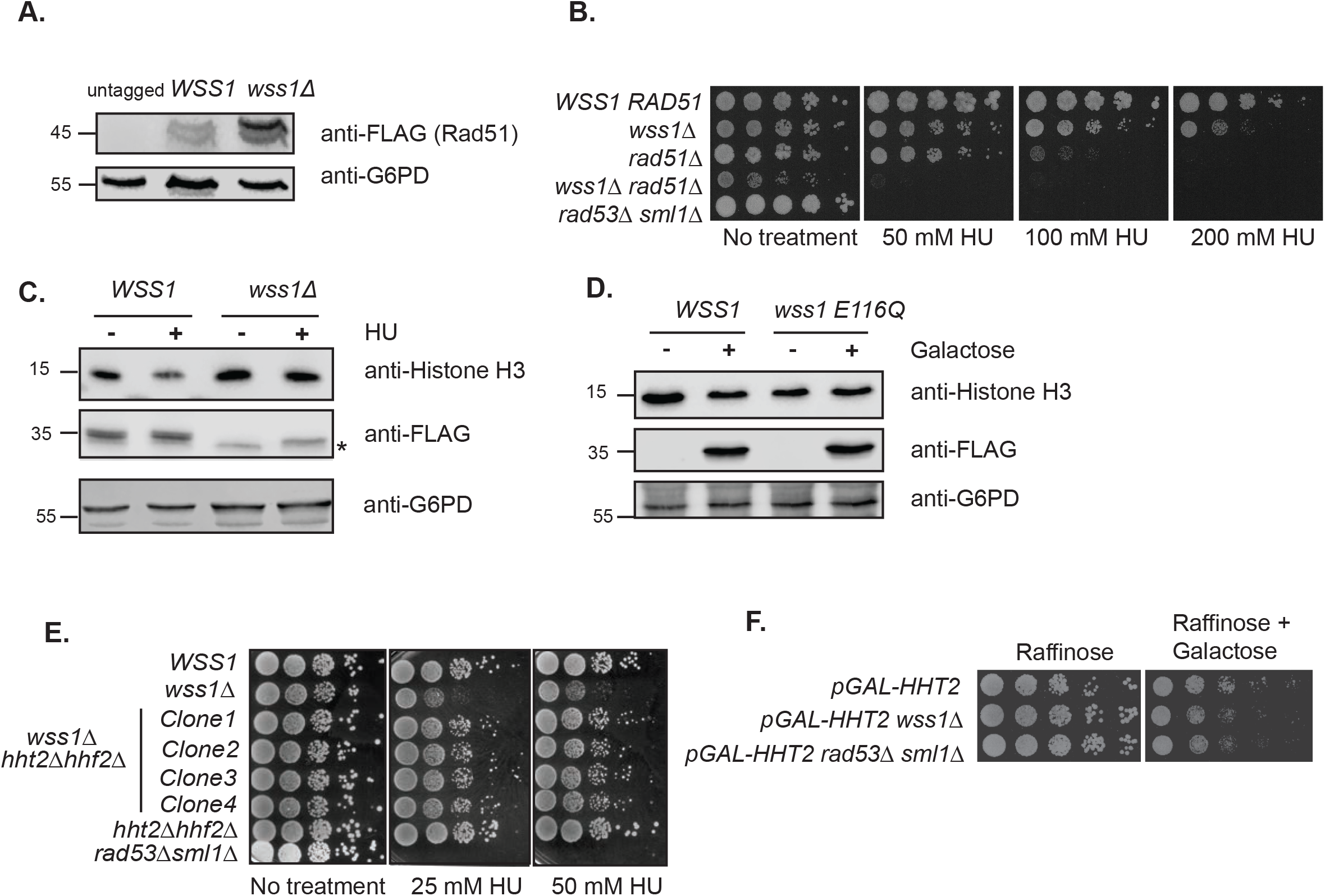
Wss1 targets histones for proteolysis. **A**) wss1Δ cells express increased Rad51 protein levels compared to wildtype cells. The endogenous Rad51 gene was 3xFLAG tagged at its C-terminus in wildtype or *wss1*Δ cells. G6PD serves as a loading control. **B**) Wss1 and Rad51 double mutants are hypersensitive to hydroxyurea. Five-fold dilutions of wildtype, *wss1*Δ, *rad51*Δ, *wss1Δrad51*Δ, and *rad53*Δ were spotted on YPD plates with or without hydroxyurea and incubated at 30°C for 3 days. **C**) Histone H3 levels decrease in hydroxyurea treated cells in a Wss1-dependent manner. Western blot analysis of endogenous histone H3 protein levels from wildtype and *wss1*Δ cells treated with or without hydroxyurea. Wss1 was 3xFLAG tagged. Asterisk indicate a non-specific band cross-reacting with the anti-Flag antibody. G6PD serves as a loading control. **D**) Decreased histone H3 protein levels depend on Wss1 catalytic activity. Western blot analysis of endogenous histone H3 protein levels from cells over-expressing wildtype or catalytic inactive Wss1 (wss1-E116Q). The expression of Flag tagged Wss1 and wss1-E116Q were placed under the control of the galactose promoter. Cells were grown in the presence of raffinose (-) or raffinose plus galactose (+) to induce expression. G6PD serves as a loading control. **E**) Reducing histone levels suppresses hydroxyurea sensitivity of *wss1*Δ cells. Five-fold dilutions of wildtype, *wss1Δ, rad53Δ, htt2Δhhf2*Δ and four independent clones of *wss1Δhtt2Δhhf2*Δ cells. Cells were spotted on YPD plates with and without hydroxyurea. Plates were incubated for 3 days at 30°C. **F**) Over-expression of histone adversely affects wss1Δ cells. Wildtype, wss1Δ or rad53Δ cells were transformed with a plasmid containing histone H3 under control of the galactose promoter. Five-fold dilutions of transformed strains were spotted on plates containing raffinose or raffinose + galactose. Plates were incubated for 3 days at 30°C.

An additional protein found in our mass spectrometry experiments, histone H3, has been previously shown to be toxic when it is expressed to high levels. We found that histone H3 and to a lesser extent linker histone H1, were enriched in hydroxyurea treated Δ*wss1* cells compared to treated wildtype cells (**Figure 2D**). We first confirmed, by western blot analysis, that histone H3 levels were regulated in a Wss1-dependent manner. An appreciable decrease in histone H3 protein levels was observed when wildtype cells are treated with hydroxyurea, when compared to *wss1*Δ cells (**Figure 3C**). Furthermore, the decrease in histone H3 protein levels depends on the proteolytic activity of Wss1 as a Wss1 catalytic-inactive mutant when over-expressed did not promote a decrease in histone H3 protein levels (**Figure 3D**).

Previous reports indicate that over-expression of histones is toxic in *rad53*Δ cells (Gunjan and Verreault, 2003; Singh et al., 2009; Singh et al., 2010). We hypothesized that in the presence of hydroxyurea, excess histones can accumulate in *wss1*Δ cells and this accumulation may cause cytotoxicity. To investigate if excess histones are causal for hydroxyurea-induced cytotoxicity in *wss1*Δ cells we sought to delete histone H3 from cells. However, the fact that histones are essential for yeast viability precluded us from deleting histone H3 from cells alone and in combination with Wss1. To circumvent this technical limitation, we reduced histone levels in *wss1*Δ and wildtype cells by deleting the *HHT2-HHF2* (H3-H4) gene pair that was shown to produce approximately seven-fold more H3-H4 transcripts than the other gene pair, *HHT1-HHF1* (Holmes and Mitchell Smith, 2001). We next generated *wss1*Δ *hht2-hhf2*Δ cells and tested four independent isolates for hydroxyurea sensitivity. Importantly and in agreement with previous reports, *hht2-hhf2*Δ cells can tolerate hydroxyurea treatment. Remarkably, deletion of the *HHT2-HHF2* gene pair suppressed the sensitivity of *wss1*Δ yeast cells to hydroxyurea suggesting that the cytotoxicity of *wss1*Δ yeast cells in the presence of hydroxyurea is due, in part, to accumulation of excess histones (**Figure 3E**). If reducing histones levels alleviates the hydroxyurea sensitivity of *wss1*Δ cells, then the over-expression of histones would adversely effect *wss1*Δ cells in the absence of hydroxyurea. To test this hypothesis, we introduced a plasmid bearing histone H3 under control of the inducible galactose promoter into wildtype cells, *wss1*Δ and *rad53*Δ cells and assessed the phenotype. A slight growth disadvantage in *wss1*Δ and *rad53*Δ cells compared to the wildtype cells was observed only when histone H3 is over-expressed with galactose (**Figure 3F**). Taken together, our results suggest that Wss1 regulate core histone levels during replication stress, which is a critical component in hydroxyurea-mediated cytotoxicity.

### Wss1 targets histones for proteolysis

To determine if Wss1 can directly exert its proteolytic function on histones, we employed an *in vitro* cleavage assay using purified recombinant proteins. Histone H3 was mixed with purified wildtype or Wss1 catalytic mutant (E116Q) in the presence of either single-stranded DNA (ssDNA) or double-stranded DNA (dsDNA). Interestingly, Wss1 cleaved histone H3 in the presence of ssDNA but not dsDNA (**Figure 4A**, lanes 8 and 9), consistent with the finding that Wss1 is activated by single stranded DNA (Balakirev et al., 2015; Stingele et al., 2014). The wss1-E116Q mutant fails to cleave histone H3 (**Figure 4A**, lanes 10 and 11), indicating that histone H3 cleavage is due to enzymatic activity of Wss1. To further accentuate this finding, we tested histone H3 cleavage in the presence of dsDNA with ssDNA over-hangs. Consistently, Wss1 could cleave histone H3 in the presence of dsDNA with ssDNA over-hangs (**Figure 4B**). To further gain molecular insight on the mechanism of histone H3 cleavage, we asked if Wss1 can act on histone H3 when all histone subunits are assembled into nucleosomes. Purified nucleosomes were incubated with wildtype Wss1 or wss1-E116Q. Interestingly, wildtype Wss1 fails to cleave histone H3 within nucleosomes suggesting that Wss1 preferentially targets histone H3 when it is bound non-specifically to single-strand DNA (**Figure 4C**). Wss1 can act on histone H3 and this effect is not specific for histone H3 alone, as wildtype Wss1 could also cleave histones H2A and H4 in the presence of ssDNA (**Figure 4D**, lanes 6 and 7). Notably, cleavage products for histone H2A by western blot were not detectable, most likely due to the loss of epitope that is recognized by the H2A antibody after Wss1 treatment. Taken together, Wss1 can directly act on core histones (H2A, H3 and H4) non-specifically bound to single-strand DNA created with hydroxyurea treatment, but not when they are incorporated into nucleosomes.

**Figure 4:**
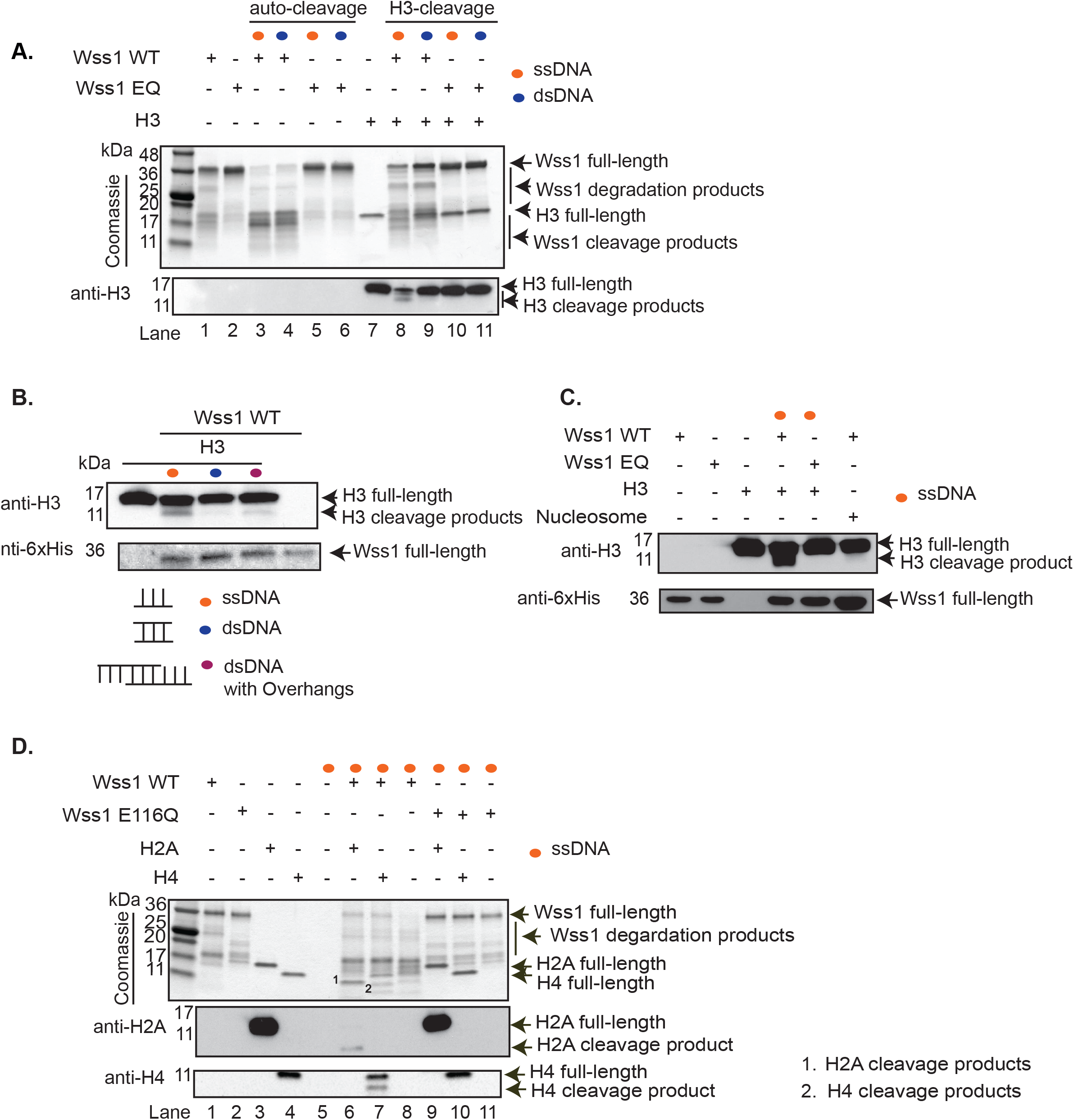
Wss1 directly targets histones for proteolysis. **A**) Wss1 cleaves histone H3 only in the presence of ssDNA but not dsDNA. Purified wild type (3.1μM) or catalytic inactive Wss1 (E116Q) (3.1μM) mutant proteins were incubated in the absence or presence of histone H3 (3.3μM) and in the absence or presence of a 73mer ssDNA or dsDNA for 2h at 30°C. Reactions were stopped by addition of 1x Laemmli sample buffer. Proteins were separated by SDS-PAGE and stained with coomassie blue to monitor Wss1 self-cleavage and histone H3 cleavage was monitored by immunoblotting with antibodies against histone H3. **B**) *In vitro* Wss1 cleaves histone H3 in the presence of dsDNA with ssDNA over-hangs. Purified wild type Wss1 (3.1μM) was incubated with histone H3 in the absence or presence of a 73mer ssDNA or dsDNA or dsDNA with ssDNA over-hangs for 2h at 30°C. Reactions were stopped by addition of 1x Laemmli sample buffer. Proteins were separated by SDS-PAGE and were further used in immunoblotting procedure to monitor histone H3 cleavage using antibodies against histone H3. **C**) Wss1 cleaves free histone H3 but not those incorporated into nucleosomes. Purified wild type Wss1 (3.1μM) was incubated with histone H3 in the presence of a 73mer ssDNA or mononucleosomes (0.42 μM) for 2h at 30°C. Reactions were stopped by addition of 1x Laemmli buffer. Proteins were separated by SDS-PAGE and were further used in immunoblotting procedure to monitor histone H3 cleavage using antibodies against histone H3. **D**) Wss1 cleaves histone H2A and H4. Purified wild type (3.1μM) or catalytic inactive Wss1 (E116Q) (3.1μM) mutant proteins were incubated in the absence or presence of histone H2A (3.3μM) or H4 (3.3μM) and in the absence or presence of 73mer ssDNA for 2h at 30°C. Reactions were stopped by addition of 1x Laemmli sample buffer. Proteins were separated by SDS-PAGE and stained with coomassie blue to monitor Wss1 self-cleavage and histone H2A and H4 cleavage was monitored by immunoblotting with antibodies against histone H2A and H4 respectively.

### Histones inhibit Wss1 self-cleavage activity

Wss1, like SPRTN, elicits self-cleavage activity in the presence of DNA (Balakirev et al., 2015; Lopez-Mosqueda et al., 2016; Stingele et al., 2014; Vaz et al., 2016). Whether Wss1 self-cleavage is a pre-requisite for substrate targeting has not been addressed. As such, we next investigated the relationship between Wss1 self-cleavage and Wss1-mediated histone cleavage. *In vitro,* Wss1 self-cleavage was readily observed in the presence of both single and double stranded DNA (**Figure 4A**, lanes 3 and 4 and **Figure 5**, lanes 1 and 2) as previously reported (Balakirev et al., 2015; Stingele et al., 2014). However, Wss1 self-cleavage is inhibited in the presence of histone H3 (**Figure 4A**, compare lanes 8 and 9 to lanes 3 and 4). In addition, this inhibitory effect on Wss1 self-cleavage, increases concomitantly with an increasing concentration of histone H3 substrate (**Figure 5A**, lanes 4-6). Wss1, although not self-cleaved, still cleaves histone H3 into distinct products (**Figure 5A**, lanes 3-6) suggesting that Wss1 self-cleavage is not a prerequisite for Wss1 activity. To gain more mechanistic insight into this inhibitory effect of histone H3 on Wss1 self-cleavage, we analyzed the kinetics of Wss1 self-cleavage in the presence or absence of histone H3. Complete Wss1 self-cleavage is achieved in forty minutes in the absence of histone H3 whereas in the presence of histone H3, Wss1 self-cleavage is significantly delayed (**Figure 5B**, **lane 6**). This raises an interesting possibility that Wss1 self-cleavage is a protease inactivating mechanism and substrates such as histones may stabilize Wss1 - preventing its premature inactivation in cells. To test this hypothesis further, we created a Wss1 mutant (wss1-RKR) that does not elicit self-cleavage in the presence of DNA. To obtain this Wss1 mutant, we first mapped the cleavage site using a rational design and substituted the amino acids at the cleavage site (**Supplemental Figure 3A**). The Wss1-RKR mutant has attenuated self-cleavage activity although it can still bind to DNA (**Supplemental Figure 3B and 3C**). Wss1-RKR was able to cleave histone H3 in the presence of single-strand DNA further supporting that Wss1 self-cleavage activity is not a requirement to cleave substrates in vitro (**Figure 5C**). Finally, *wss1*Δ cells were reconstituted with the *wss1-RKR* mutant allele. We found that cells expressing the wss1-RKR mutant allele can tolerate hydroxyurea as wildtype cells (**Figure 5D**). Taken together, our results would suggest that Wss1 substrates can regulate Wss1 self-cleavage.

**Figure 5:**
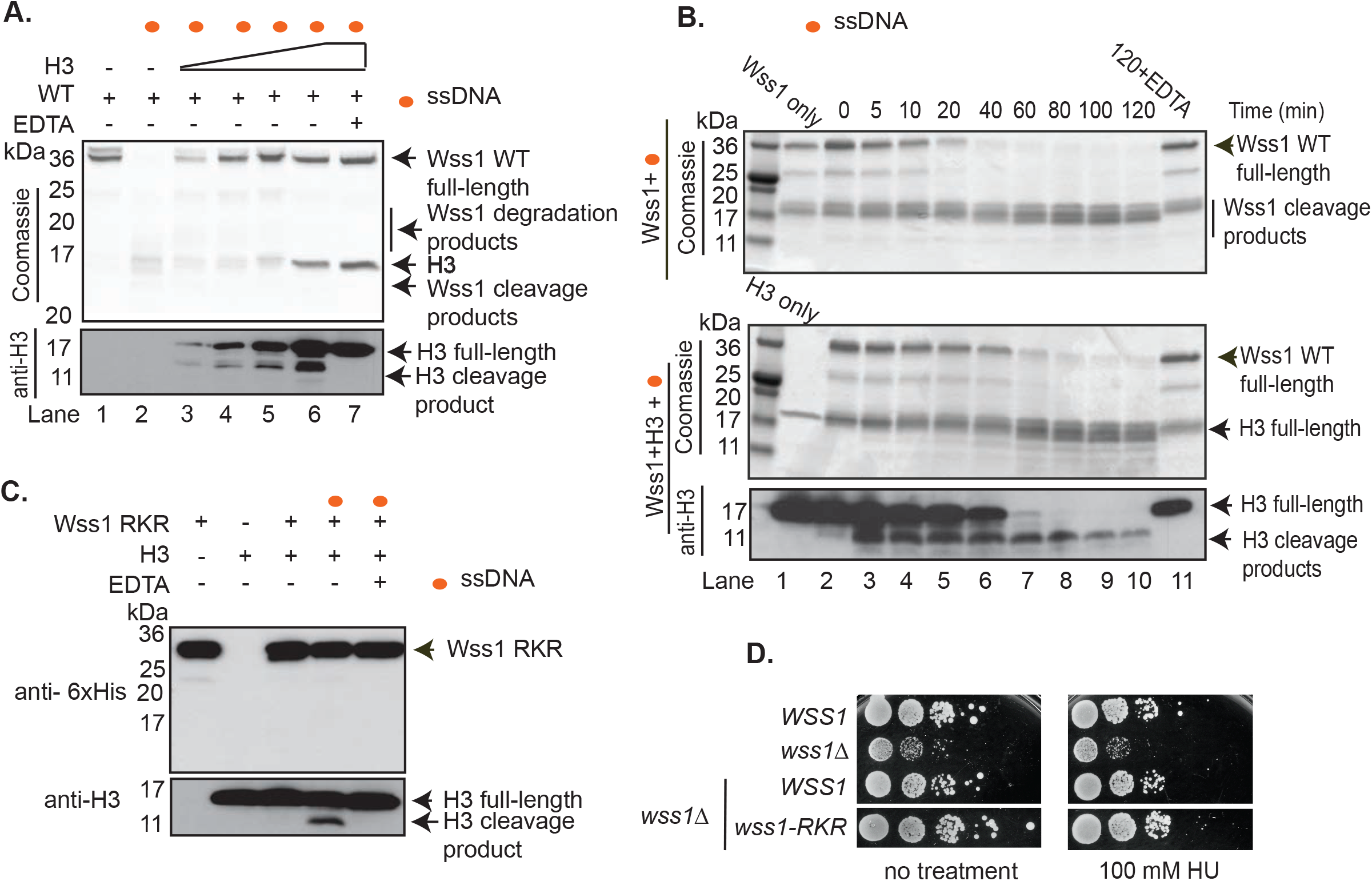
Wss1 self-cleavage is not a requirement for proteolytic activity. **A**) Titration of histone H3 in Wss1 auto-cleavage reactions *in vitro*. Purified wildtype Wss1 (3.1μM) and varying concentrations of histone H3 (3.3, 6.6, 13.2, 26.4 μM) were incubated in the presence of a 39mer ssDNA (10μM) for 2h at 30°C. A control reaction without histone H3 or ssDNA was used in the experiment. Reactions were stopped by addition of 1x Laemmli sample buffer. Proteins were separated by SDS-PAGE and stained with coomassie blue to monitor Wss1 self-cleavage and histone H3 cleavage was monitored by immunoblotting with antibodies against histone H3. **B**) The presence of H3 delays Wss1 self-cleavage. Purified wildtype Wss1 (3.1μM) was incubated with or without histone H3 (3.3 μM) in the absence or presence of ssDNA (10μM) for 2h at 30°C and samples were taken at indicated time points. Proteins were resolved by SDS-PAGE and stained with Coomassie blue to monitor the auto-cleavage of Wss1 and H3 cleavage was monitored by immunoblotting with antibodies against histone H3. **C**) A self-cleavage Wss1 mutant has proteolytic activity. Purified Wss1 RKR (3.1 μM) was incubated with histone H3 (3.3 μM) in the absence or presence of ssDNA (10μM) for 2h at 30°C. Reactions were stopped by addition of 1x Laemmli sample buffer. Proteins were resolved by SDS-PAGE and used for immunoblotting to monitor the presence of Wss1-RKR or histone H3 cleavage using antibodies against 6xHis-tagged Wss1 RKR and H3 antibodies respectively. D) Wss1-RKR can rescue the hydroxyurea-sensitivity of *wss1*Δ cells. Five-fold dilutions of wild type or *wss1*Δ cells reconstituted with either wild type or wss1 RKR mutant alleles, spotted on plates in the absence or presence of hydroxyurea (100 mM) and incubated at 30°C. Plates were imaged after 3 days.

## Discussion

We have utilized the sensitivity of *wss1*Δ cells to hydroxyurea to identify histones as a novel class of Wss1 substrates, which are distinct from covalent DPC substrates. Histones are known to form insoluble protein-DNA aggregates when expressed in excess (Clark and Kimura, 1990). In cells histone abundance is carefully regulated at many levels (Osley, 1991). During the S-phase, histone chaperones sequester histones and deposit them into nascent DNA. Wss1 cannot degrade histones assembled in nucleosomes but can degrade histones that are binding to single-stranded DNA. When cells are exposed to hydroxyurea, large tracks of single stranded DNA are exposed at DNA replication forks and newly synthesized histones cannot be assembled into nucleosomes. As such, the histones will bind non-specifically to any nucleic acid and such high affinity/avidity binding of histones to DNA could interfere with DNA metabolism. Fob1 is one notable example of a protein that strongly binds to DNA in budding yeast. Fob1 binding to DNA is sufficient for blocking DNA replication forks at the rDNA locus(Brewer and Fangman, 1988; Kobayashi and Horiuchi, 1996). It has been previously shown that histone abundance is cytotoxic by saturating binding to histone modifying enzymes as well as non-specific binding to DNA and RNA (Singh et al., 2010).

Hydroxyurea treatment is more toxic to *wss1*Δ cells overexpressing Wss1 catalytic dead mutants (E116Q) than to *wss1*Δ cells (**Figure 1F**). This toxicity could be explained by Wss1-E116Q binding to protein substrates but not cleave them and thereby form higher complexity enzyme-substrate complexes on DNA, but alternative models are also possible. Reducing histone dosage by deleting the H3-H4 gene pair (*HHT2-HHF2*) suppressed the hydroxyurea sensitivity in *wss1*Δ cells, suggesting that cytotoxicity of *wss1*Δ yeast cells in the presence of hydroxyurea is due, in part, to non-specific binding of histones to DNA. It is possible that deleting *HHT2-HHF2* from cells strongly confers resistance to hydroxyurea irrespective of Wss1 function. However, deletion of histones does not suppress hydroxyurea sensitivity in every case. For instance, the exoribonuclease Xrn1, when deleted, renders cells hypersensitive to hydroxurea and this sensitivity is not suppressed when histone are deleted in an *xrn1* mutant background(Lao et al., 2018).

A direct link between histone proteolysis and Wss1 during hydroxyurea treatment was obtained by *in vitro* cleavage assays showing that Wss1 could degrade histone H2A, H3 and H4. Importantly, Wss1 could target histones only in the presence of single strand DNA, a finding that is consistent with vast single strand DNA exposed with hydroxyurea treatment. Excess histones non-specifically binding to nucleic acids is likely to have adverse effects in cells due to interference with polymerases important for DNA replication, gene transcription and translation, as all these cellular processes have DNA as an initial common template (Edenberg et al., 2014). In the case of *wss1*Δ cells during replication stress, we speculate that histones pose a problem to gene transcription machinery rather than to DNA replication *per se*, as we did not detect problems with global DNA replication under our experimental conditions. It is possible that in *wss1*Δ cells, other pathways cooperate to preserve DNA replication fork progression. Indeed, the histone H3 chaperone, Asf1, was shown to buffer excess histones during replication stress (Groth et al., 2005). In addition, Rad53 mediates a phosphorylation-dependent degradation of excess histones and this mechanism is also likely to function during replication stress (Gunjan and Verreault, 2003; Singh et al., 2009). More recently, Ddi1 was implicated to function in parallel to Wss1 for providing replication stress tolerance(Serbyn, 2019; Svoboda et al., 2019). Taken together, cells employ multiple pathways to counteract excess histones during replication stress. Wss1 dependent degradation of histones is one way yeast cells are able to tolerate replication stress in order to maintain the integrity of the genome.

## Experimental procedures

### Yeast Strains

W303a yeast cells were used to generated *Δ*wss1 yeast strain by targeted gene deletion using standard yeast methods (yeast gene deletion plasmids: pAG32-hphMx, pAG25-CloNAT, pFA6a-TRP1). To generate *Δ*wss1 yeast strains expressing 2xFLAG-Wss1 wildtype or various mutant alleles or 2xFLAG-SPRTN wildtype or catalytic mutant allele under the control of the galactose inducible promoter, Wss1 was cloned into pDONR223 (Invitrogen) using the Gateway cloning system and subsequently cloned into pAG-306GAL-ccdB (pAG-306GAL-ccdB was a gift from Susan Lindquist Addgene plasmid #14139). The plasmid was linearized at the URA3 loci and transformed into yeast strains. PCR cassette based C-terminal tagging was employed to generate Rad52-GFP and subsequent yeast strains (plasmid used for PCR cassette:pFA6a-link-yEGFP-KanMX6)

### Yeast spot assays

Yeast strains were grown overnight at 30°C with shaking for yeast spot dilution assays. Next day, 0.2 OD (approximately 2 × 10^5^ cells) of cells including ten- or five-fold dilutions prepared in sterile ddH_2_O. Cells were platted as spots in either YPD (Yeast extract, peptone and Dextrose) or YPG (Yeast extract, peptone and galactose) or YPD or YPG plates containing drugs hydroxyurea or CPT or H_2_O_2_. Plates were incubated at 30°C and photographs of the plates were taken as indicated in the figure legends.

### Mass Spectrometry based analysis of yeast whole cell extracts

Protein from yeast whole cell extracts were separated by 1D PAGE, each gel-lane was cut in 12 pieces and subjected to in-gel trypsin digestion. Tryptic peptides were desalted by StageTip cleanup and analyzed by LC-MS/MS. In brief, the peptides were separated by C18 reversed phase chromatography with an Easy nLC 1200 (ThermoFisher) coupled to a Q Exactive HF mass spectrometer (ThermoFisher). For each protein identification and label-free quantification, spectra were extracted and searched against the Uniprot *Saccharomyces cerevisiae* database with Maxquant.

### Expression and Purification of Wss1 wildtype and mutant alleles

Wss1 wild type or mutant alleles was cloned in to pET15 b vector with a N-terminal 6x Histidine tag and a 3C-protease (in-house produced) site using restriction sites (Nde1/Xho1) by DNA restriction method. Wss1 was expressed in Rosetta *E. coli* cells. Expression was induced with 0.25mM IPTG overnight at 18°C. Cells were harvested by centrifugation and lysed in lysis buffer (50mM Tris pH 7.5, 1M NaCl, 10% glycerol, 10mM imidazole) and performed Affinity based batch purification using TALON affinity resin. The protein was eluted using elution buffer containing Imidazole (50mM Tris pH7.5, 200mM NaCl, 200mM Imidazole). A final round of size-exclusion chromatography was performed using Superdex 75 16/60 column in the buffer 20mM Tris pH 7.5, 300mM NaCl. Fractions containing monomeric form of wild type or mutant proteins were pooled and were further concentrated using centrifugal filter (10kD cutoff) to a final concentration of 2 mg/ml and flash frozen at −80°C for further use.

### Wss1 self-cleavage assays

Purified Wss1 wildtype (3.1 μM) or various mutant alleles (3 μM) were incubated with ssDNA (Sigma) (5’GCGCGCCCATTGATACTAAATTCAAGGATGACTTATTTC3’) (10μM) at 30°C for 2 hrs. EDTA (2mM) was used to inhibit the reactions. Reactions were stopped by adding 1x Laemmli buffer and samples were boiled at 95°C for 10 min. Further, proteins were resolved using SDS-PAGE and further transferred to a nitro-cellulose membrane for Western blot analysis.

### Histone titration in the presence of Wss1 and ssDNA

Purified Wss1 wild type (3.1 μM) was mixed with purified histone H3 (NEB) (3.3 μM, 6.6 μM, 13.2 μM and 26.4 μM) in the presence of ssDNA (10μM). Reactions were incubated at 30°C for 2 hrs and stopped by addition of 1x Laemmli buffer and boiled at 95°C for 10 min. Further, samples were analysed using SDS-PAGE analysis. Auto-cleavage of Wss1 was monitored by staining the protein using Coomassie. Histone H3 cleavage was monitored using anti-H3 antibody using western blot analysis.

### Kinetics of Wss1 auto-cleavage and histone H3 cleavage

Purified Wss1 wild type (3.1 μM) was mixed with ssDNA (10μM) in the presence or absence of purified histone H3 (3.3 μM). Reactions were incubated at 30°C for 2 hrs. Samples were collected at indicated time points and stopped by addition of 1x Laemmli buffer and boiled at 95°C for 10 min. Further, samples were analysed using SDS-PAGE analysis. Auto-cleavage of Wss1 was monitored by staining the protein using Coomassie. Histone H3 cleavage was monitored using anti-H3 antibody using western blot analysis.

### EMSA (Electrophoretic mobility shift assay)

Purified proteins in indicated concentrations were incubated with fluorescently labelled ssODN (Sigma) (0.25μM-6-FAM-39mer-5’GCGCGCCCATTGATACTAAATTCAAGGATGACTTATTTC3’) in the reaction buffer (10mM Tris 7.5, 0.2mM DTT, 5μM ZnSo4) and incubated at 20°C for 30 min. Protein-DNA complexes were separated on 1.5% Agarose gel and visualised by FUSION-SL imager (Vilber)

### Flow cytometry (FACS)

For flow cytometry analysis of yeast cells, yeast cells were collected at respective indicated time points and fixed them immediately with 70% EtOH in flow cytometry tubes. Next, cells were sonicated (20% amplitude, 3 sec) to separate cell clumps. Cells were centrifuged at 300 rpm for 5 min and ethanol was aspirated out. Later, cells were re-suspended in 1ml of RNAse buffer (50mM Tris pH8.0 and 15mM NaCl) with RNAse (0.25 mg/mL) and incubated at 50°C for 1 hour. Further, proteinase K (0.125 mg/mL) was added to the buffer and incubated for additional 1 hr at 50°C. Finally, propidium Iodide at a dilution of 1:1000 was added and vortexed to have final sample preparation. Readings were recorded on BD FACS™CONTO II and FACS curves were analysed using FlowJo software.

### Preparation of yeast whole cell extracts for western blot analysis

TCA precipitation method was employed to obtain the protein extracts used in western blot analysis. For yeast protein extracts, 5-10 O.D cells were harvested by centrifugation were re-suspended in 500μl of TCA lysis buffer (10mM Tris pH 8.0, 20% TCA, 25mM NH_4_CH_3_CO2) to precipitate the proteins and centrifuged to remove the supernatant. Precipitated protein pellets were washed with 70% Acetone and re-suspended in 75 μl buffer containing 10mM Tris pH 11.0 and 3% SDS and 75 μl of 1xLaemellie buffer was added to make a final volume of 150 μl and samples were boiled at 95°C for 10 min.

## Acknowledgements

We are grateful to Susan Lindquist for providing plasmids, Akash Gunjan for providing the histone reduced yeast strain (*Δhht2-hhf2*), Helle Ulrich for providing Rad53 mutant yeast strains. We thank Dikic lab members for their continued support and constructive discussions. Jaime Lopez-Mosqueda was supported by a long-term post-doctoral fellowship from the Human Frontiers Science Program and South Dakota Agricultural Experiment Station. Daniel Sam is supported by South Dakota Agricultural Experiment Station funds. This work was supported by grants from the DFG (SFB1177), the Cluster of Excellence “Macromolecular Complexes” of the Goethe University Frankfurt (EXC115), LOEWE grant Ub-Net and LOEWE Centrum for Gene and Cell Therapy Frankfurt.

## Author contributions

KM, JLM and ID conceived the project. KM and JLM designed all the experiments and ID supervised the project. KM constructed *Δwss1* based yeast strains and performed all the experiments including yeast and *in vitro* enzymatic assays. FB assisted with mass spectrometry based methods and data analysis. DKS and ET created yeast strains and performed experiments under supervision from JLM. SP assisted with protease dependent degradation of histone H3 in cells. KM, JLM, and ID wrote the manuscript.

---

**Supplementa1 Figure 1:**
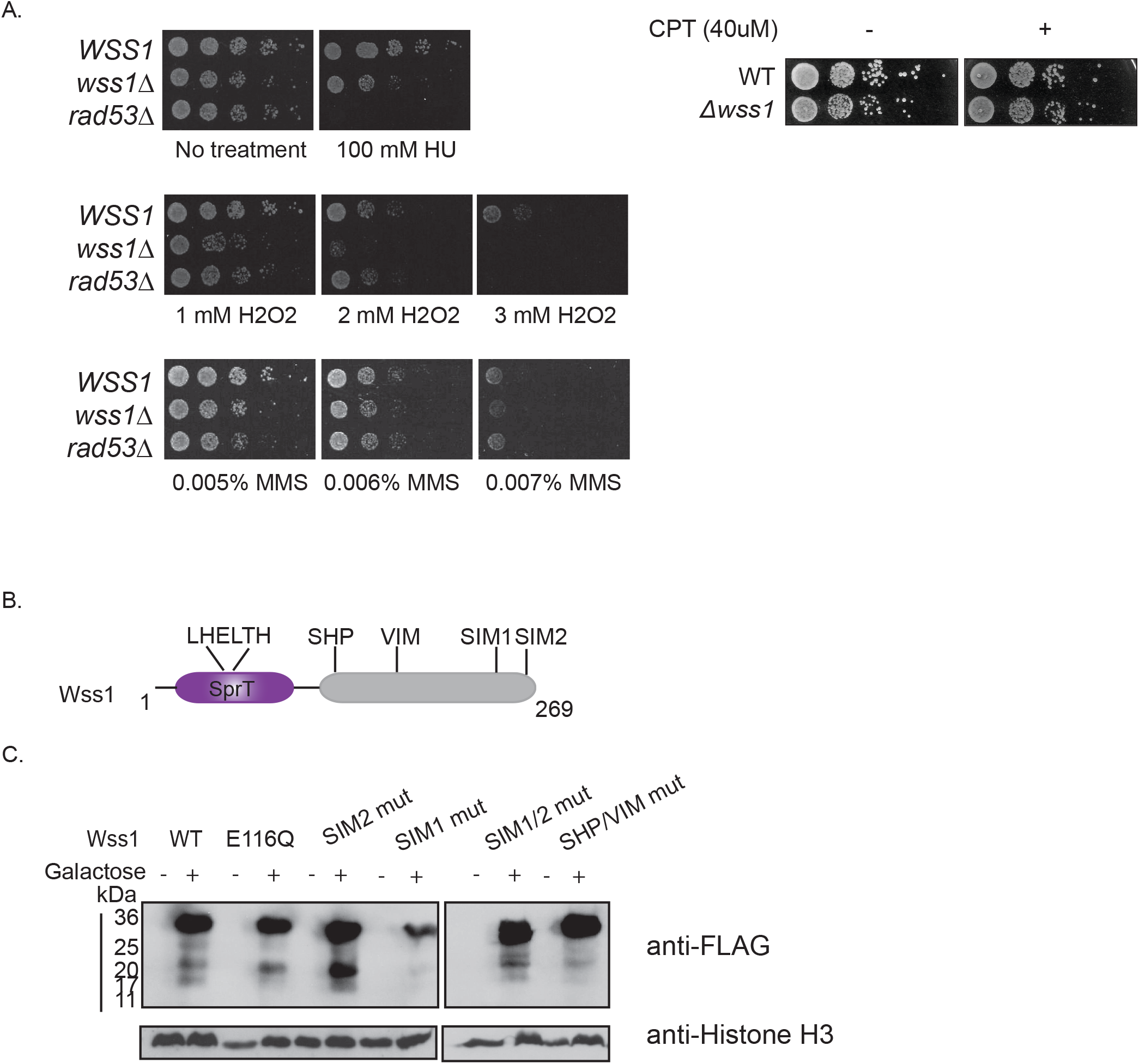
Wss1 deficient cells are sensitive to hydroxyurea and hydrogen peroxide. **A**) Assessment of sensitivity of *wss1*Δ cells towards various genotoxic agents. Ten-fold dilutions of wild type or *wss1*Δ yeast cells were spotted on plates in the absence or presence of H2O2 (3mM) or Hydroxyurea (150mM) or CPT (40μM). The plates were incubated at 30°C and imaged after 3 days. **B**) Domain organization of Wss1. **C**) Expression confirmation analysis of FLAG-tagged Wss1 and various mutant alleles in Δwss1 cells. Cell extracts were prepared by TCA precipitation method and immuno-blotted using anti-FLAG antibody (Sigma) or histone H3 antibody (Abcam).

**Supplemental Figure 2.**
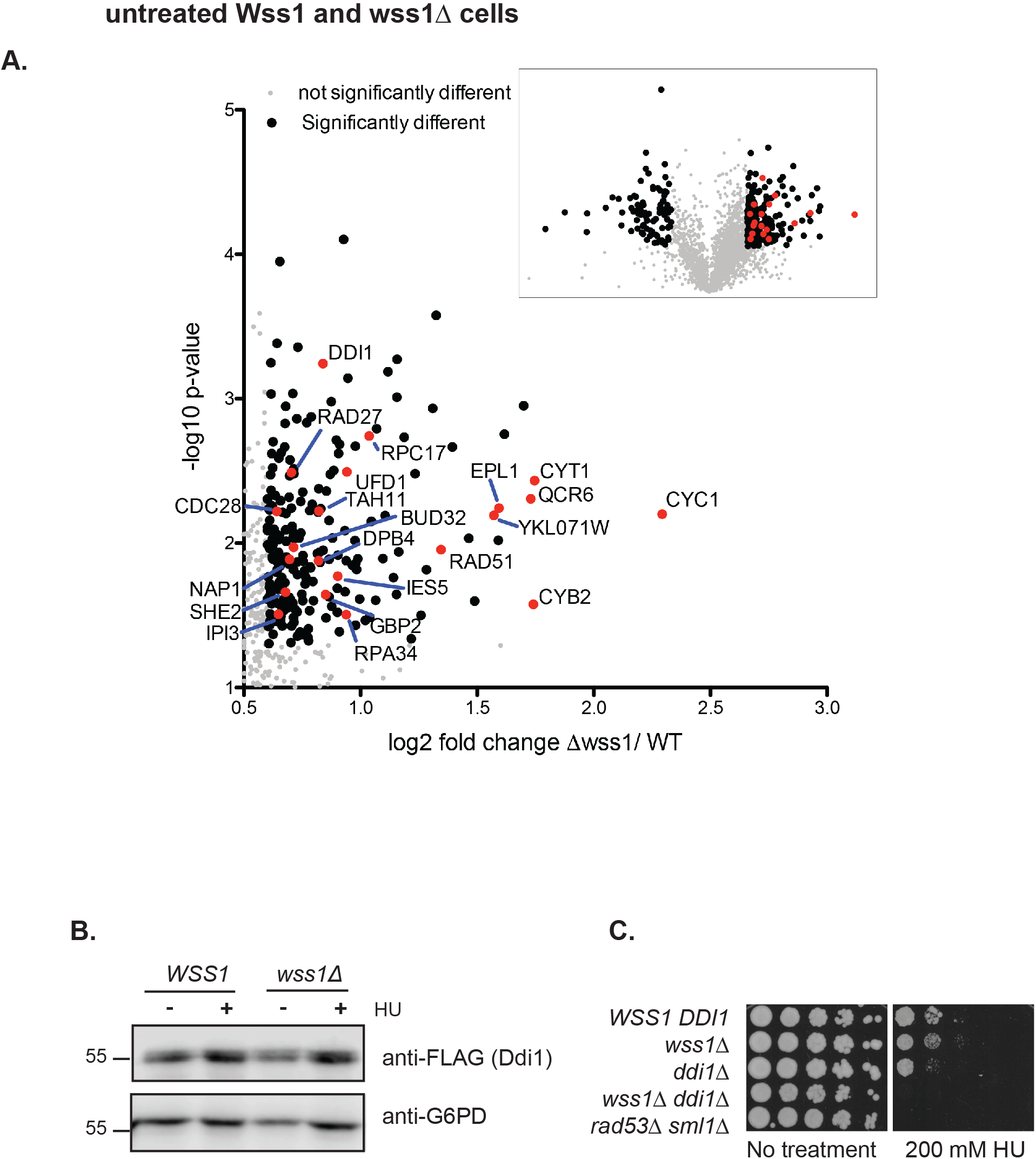
Mass spectrometry-based proteome quantification in untreated cells. (**A**) Volcano plots representing SILAC based quantification of peptides from untreated Wild type and *wss1*Δ cells. Inset represents the results of three independent mass spectrometry quantifications of proteome changes in wildtype (left) and *wss1*Δ cells (right). Gray circles represent proteins that were not statistically enriched. Black circles represent proteins that were significantly enriched *t-test p<0.5.* **B**) Hydroxyurea treated *wss1*Δ cells express increased Ddi1 protein levels compared to untreated cells. The endogenous Ddi1 gene was 3xFLAG tagged at its C-terminus in wildtype or *wss1*Δ cells. G6PD serves as a loading control. B) Wss1 and Ddi1 double mutants are hypersensitive to hydroxyurea. Five-fold dilutions of wildtype, *wss1*Δ, *ddi1*Δ, *wss1Δddi1*Δ, and *rad53*Δ were spotted on YPD plates with or without hydroxyurea and incubated at 30°C for 3 days.

**Supplemental Figure 3:**
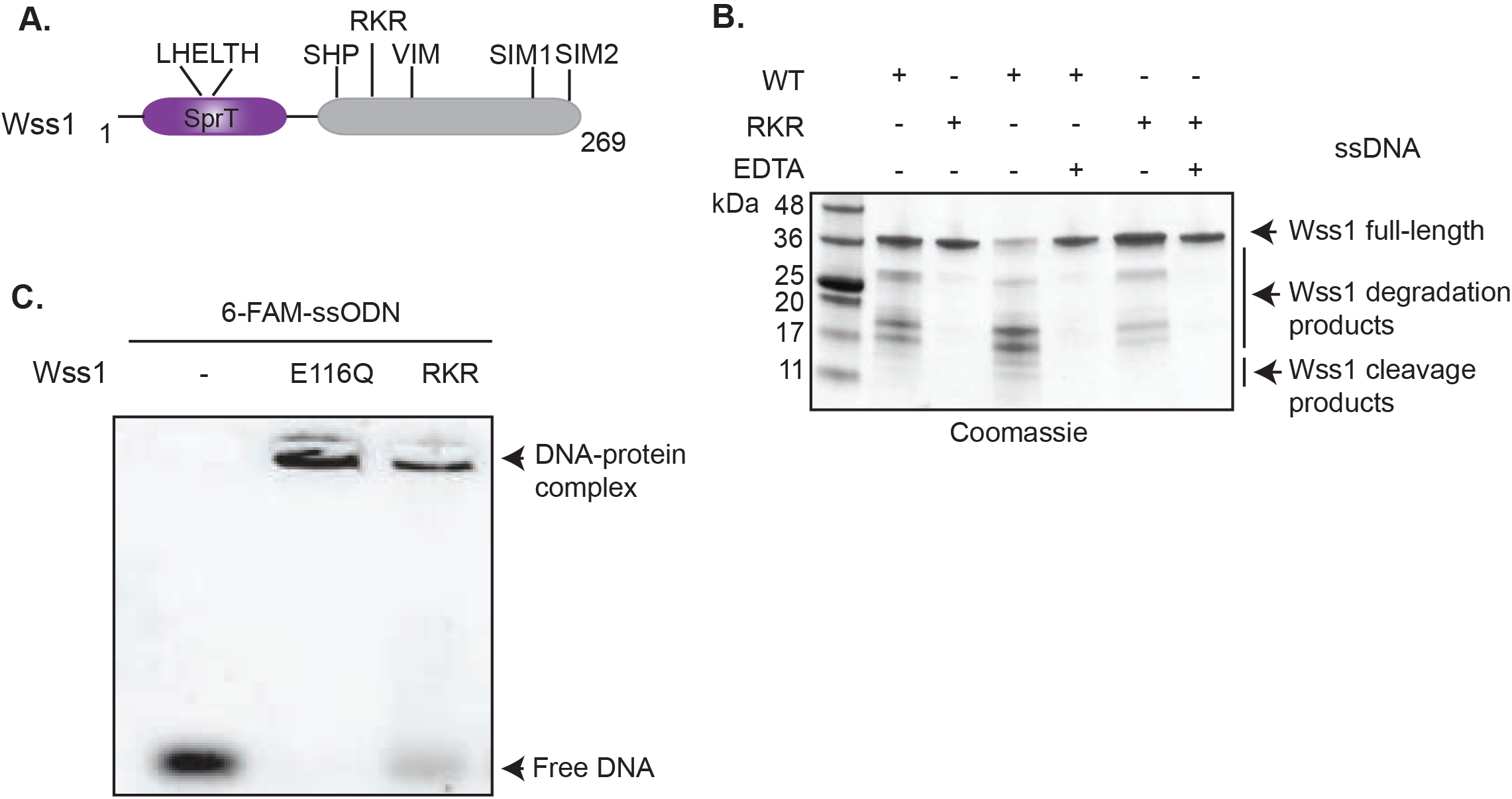
Characterization of Wss1-RKR mutant. (**A**) Depiction of RKR site in the domain organization of Wss1. **B**) Wss1-RKR mutant is self-cleavage deficient. Purified wild type (3.1μM) or Wss1-RKR (3.1μM) mutant proteins were incubated in the absence or presence of a 73mer ssDNA and in the absence or presence of EDTA (2mM). Reactions were stopped by addition of 1x Laemmli buffer. Proteins were separated by SDS-PAGE and stained with Coomassie blue. **C**) Wss1-RKR binds to single stranded DNA. EMSA analysis of Wss1 full-length (E116Q) and Wss1-RKR in the presence of a fluorescently labelled 6’FAM-ssODN.

